# Ultralow frequency vaso-oscillations in human cerebral arteries are independent from Mayer waves

**DOI:** 10.64898/2026.06.09.731141

**Authors:** Aiman Alzetani, Jacob Duckworth, Anthony A Birch, David M Simpson, David Kleinfeld, Roxana O Carare

## Abstract

This study tests the hypothesis that vasomotion, an ∼ 0.1 Hz oscillation in arteriole diameter, is generated by intrinsic oscillations within the arterioles that perfuse the brain, and not by external drive from systemic blood pressure oscillations (Mayer waves). During cardio-pulmonary bypass that transiently eliminated systemic blood pressure oscillations in 14 patients, we observed that vasomotor oscillations persist with normal amplitudes and frequencies over the one- to three-hour time course of surgery. In contrast, ∼ 0.1 Hz oscillations in peripheral blood pressure were predominantly absent. This implies that cerebral arterioles generate their own rhythmic vaso-dynamics, although we cannot discount that vasomotion can phase-lock with ∼ 0.1 Hz systemic physiological rhythms in the awake, healthy state. We discuss the impact of this finding on the role of vasomotion in modulating the perfusion of blood and the transport of interstitial fluid in the brain.

## Introduction

Perfusion to the brain is maintained during variations in peripheral blood pressure through autoregulatory changes in the diameter of major and minor feeding arteries to the organ^1^. Dynamic cerebral autoregulation acts within seconds on vascular smooth muscle cells (vSMCs), and consequently, slow peripheral blood pressure oscillations can entrain oscillations in cerebral artery diameter. This occurs during both experimentally forced blood pressure changes^2^ as well as during spontaneous Mayer wave events^3^, which are observed during resting conditions^3, 4^. Arteries throughout the body also undergo ∼ 0.1 Hz vasomotion driven by intrinsic vSMC activity^5, 6^. Vasomotion has been observed in various tissues and species, including intraoperatively in pial arterioles of awake humans^4, 7^. However, dissociating intrinsic vasomotion from sources of external drive is notoriously difficult. In the brain, cerebral arterioles respond in diameter to adrenergic or cholinergic input from locus coeruleus/nucleus basalis of Meynert^8, 9^, can be entrained by ultraslow ∼ 0.1 Hz variations in cortical neural activity^10^, and can phase-lock with systemic oscillations near ∼ 0.1 Hz^11-13^. In humans, the Mayer wave frequency overlaps spectrally with vasomotion, which further masks the origin of observed ∼ 0.1 Hz vaso-oscillations^3^.

Vasomotion has recently gained attention for its likely contribution in intramural periarterial drainage (IPAD)^14^, which facilitates the removal of toxic metabolites, including amyloid-β (Aβ) peptides, that are implicated in Alzheimer’s disease. IPAD relies on ∼ 0.1 Hz vasomotion for the efficient clearance of Aβ and other metabolic waste^14, 15^. The mechanism for net flow in the face of oscillatory fluidic motion is experimentally unresolved^16^, although vasomotion is predicted to induce a net volume flow rate^14^owing to the asymmetry of the arteriolar oscillations during functional hyperemia^17, 18^, the potential for peristalsis^14, 19^, and asymmetric flow through poroelastic tissue^20^. Lastly, in aging and Alzheimer’s disease, cerebrovascular function is often disrupted, leading to reduced waste clearance. the accumulation of Aβ plaques, and exacerbated neurodegeneration and cognitive decline^21^.

For both reasons of fundamental physiology and biomedicine, it is crucial to identify the driving force for cerebral vasomotion. Oscillations in cerebral perfusion pressure, termed Mayer waves, are driven by feedback between sympathetic nerve activity and changes in peripheral vascular bed resistance^3^. In rodents, the Mayer wave frequency is approximately 0.4 Hz^22^, whereas in humans the frequency is near 0.1 Hz and thus overlaps spectrally with vasomotion and obfuscates the origin of ∼ 0.1 Hz hemodynamic fluctuations. To disambiguate vasomotion from Mayer waves, we sought to measure cerebral blood flow in the absence of Mayer waves by artificially stabilizing systemic blood pressure^23^.

One of the most significant advances in the management of heart diseases came from the introduction of cardiopulmonary bypass (CPB)^24^. A machine is connected to the heart to divert blood away from it and the lungs while operating on an arrested “motionless” heart, typically to replace diseased valves or correct congenital malformations. The CPB machine supports the systemic circulation and gas exchange for the whole body, including the brain. The flow, speed, pressure, volume, temperature, pH, and blood gas concentrations can be manipulated to maintain “normal” physiological homeostasis as much as possible. CPB also allows us to create abnormal states, such as cooling the blood to reduce brain metabolic activity, reducing the cerebral oxygen requirements and allowing reduced blood flow to the body for short spells while facilitating a complex cardiovascular reconstructive procedure^25^. Since CPB removes cardiac variability and artificially stabilizes aortic pressure, it provides the ideal condition to test if cerebral blood flow oscillations depend on systemic Mayer waves.

## Material and Methods

### Patients and procedures

A protocol driven observational study was set up under ethical approval - Integrated Research Application System (IRAS) (Number 264726); complete statements on ethical considerations are given at the end matter. Participants were recruited from patients who were admitted for cardiothoracic surgical procedures. All patients had general anesthesia with routine monitoring of radial arterial blood pressure (ABP), central venous pressure (CVP), arterial blood gases, central and peripheral temperatures, heart rate, pulse oximetry and ECG. These were connected to a monitor for continuous data display and recording (MetaVision®, iMDsoft, Tel Aviv, Israel). The CPB system was LivaNova Sorin Stockert S5, widely used in cardiopulmonary bypass. This system contains a peristaltic pump that was set to deliver a constant flow, as opposed to a pulsatile flow or a constant pressure. Nonetheless, the frequency of the pump in the CPB machine is BPM/60 Hz, where we used a BPM of 120/minute, so the pump head generated pulsatile oscillations at 2 Hz. There is no internal mechanism in the pump that generates a 0.1 Hz signal. A final point is that the ventilator, which runs at 0.2 Hz, is only on before and after, but never during, CPB. Thus the ventilator cannot be a source of 0.1 Hz during CPB.

### TCD recording

There was additional monitoring of the middle cerebral artery using a transcranial Doppler ultrasound (TCD) Dopplerbox (DWL GmbH, Singen, Germany). This was connected via an analogue to digital converter, to a PC which was streaming digital outputs from the clinical monitors. The TCD and CPB signals were simultaneously recorded at 240 Hz using ICM+ software (ICM+, Cambridge Enterprise, Cambridge, UK, (http://www.neurosurg.cam.ac.uk/icmplus).

After the patient had been transferred to theatre, and the anesthetic team were satisfied with their lines and monitoring, surgery began, and the transcranial Doppler probe was adjusted to obtain an optimal signal from either left or right middle cerebral artery. Recording started as soon as a good signal had been obtained. It was maintained throughout the operation until wound closure. The arterial blood pressure varied around a mean of 65 mmHg. From the full length of TCD recording we manually discarded times when the signal was either completely absent or corrupted by too much motion or diathermy artifact (**Figure 1**). We considered a clear TCD signal of 5 minutes in length as “good” data for analysis. All good-quality data sections of 5 minutes duration were used in the analysis. One trial of 3.6 minutes and one of 4 minutes were also included. Data sections were identified according to the stage of the operation as before cardiac bypass (pre-CPB), on bypass (on-CPB) or post-bypass (post-CPB).

**Figure 1:**
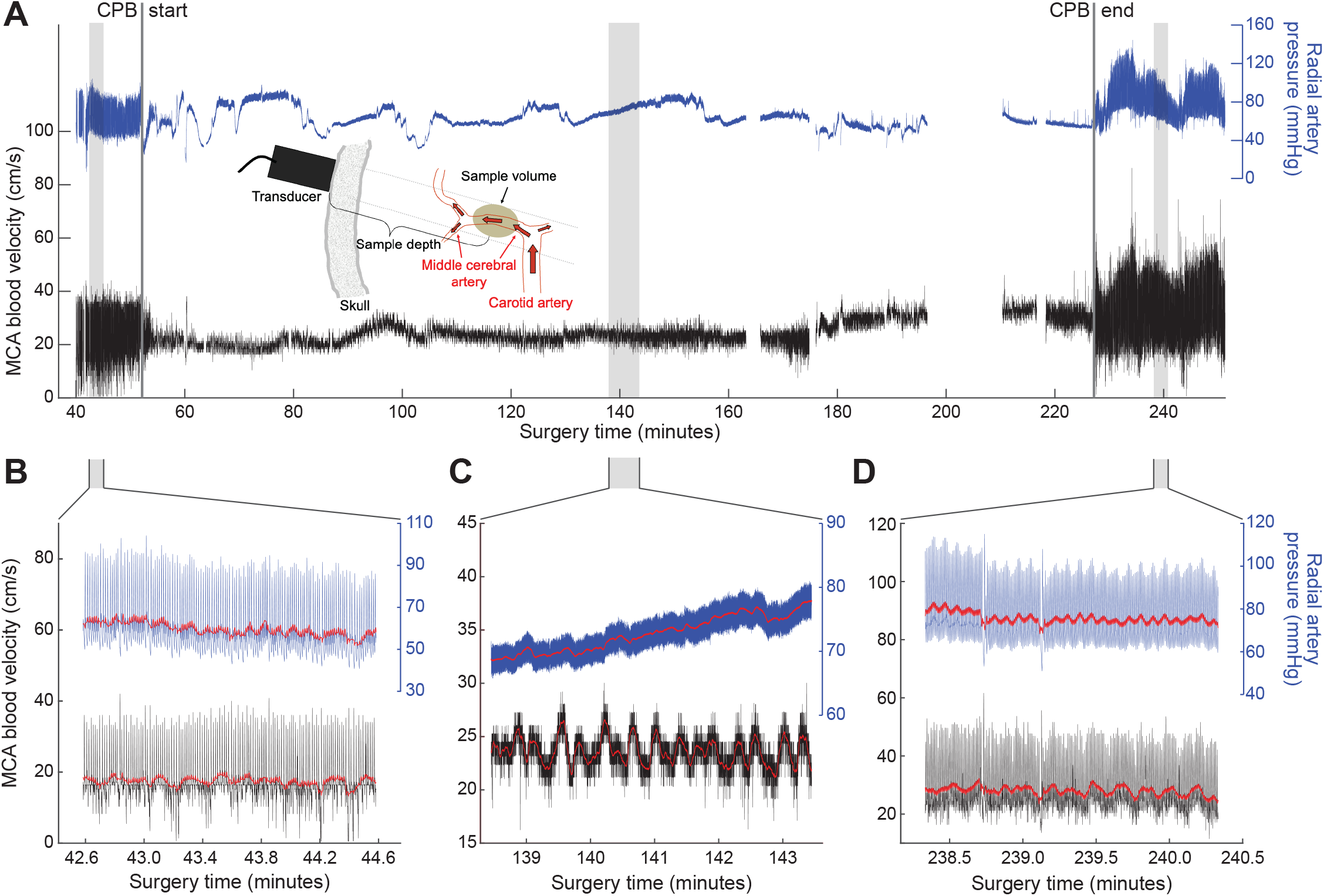
Full time traces of a patient undergoing CPB. **A**, MCA Blood Velocity trace in cm/s (Black), radial artery blood pressure trace in mmHg (Blue). **B-D**, Three segments of “good data” for analysis are marked and expanded: before (**B**), during (**C**), and after (**D**) CPB, with two-second moving mean traces overlaid (Red). Inset: a sketch of transtemporal positioning of a TCD probe to detect flow from the MCA. Abbreviations: TCD, transcranial Doppler, MCA, middle cerebral artery, CPB, cardiopulmonary bypass.

### Data analysis

This applies to the data of **Figure 2**. Outlier removal was performed on each time series by detecting and replacing discontinuities of more than two standard deviations from the mean and lasting less than 200 ms, with a linear interpolation of the surrounding data. The best fit linear trend was then removed before spectral analyses. Multitaper spectral analysis was performed in MATLAB using Slepian tapers^26^. Power spectral density (PSD) and coherence estimates were calculated using a half-bandwidth, denoted Δf, of Δf = 0.02 Hz and 2xTxΔf - 1 orthonormal tapers, where T is the trial length in seconds.

**Figure 2:**
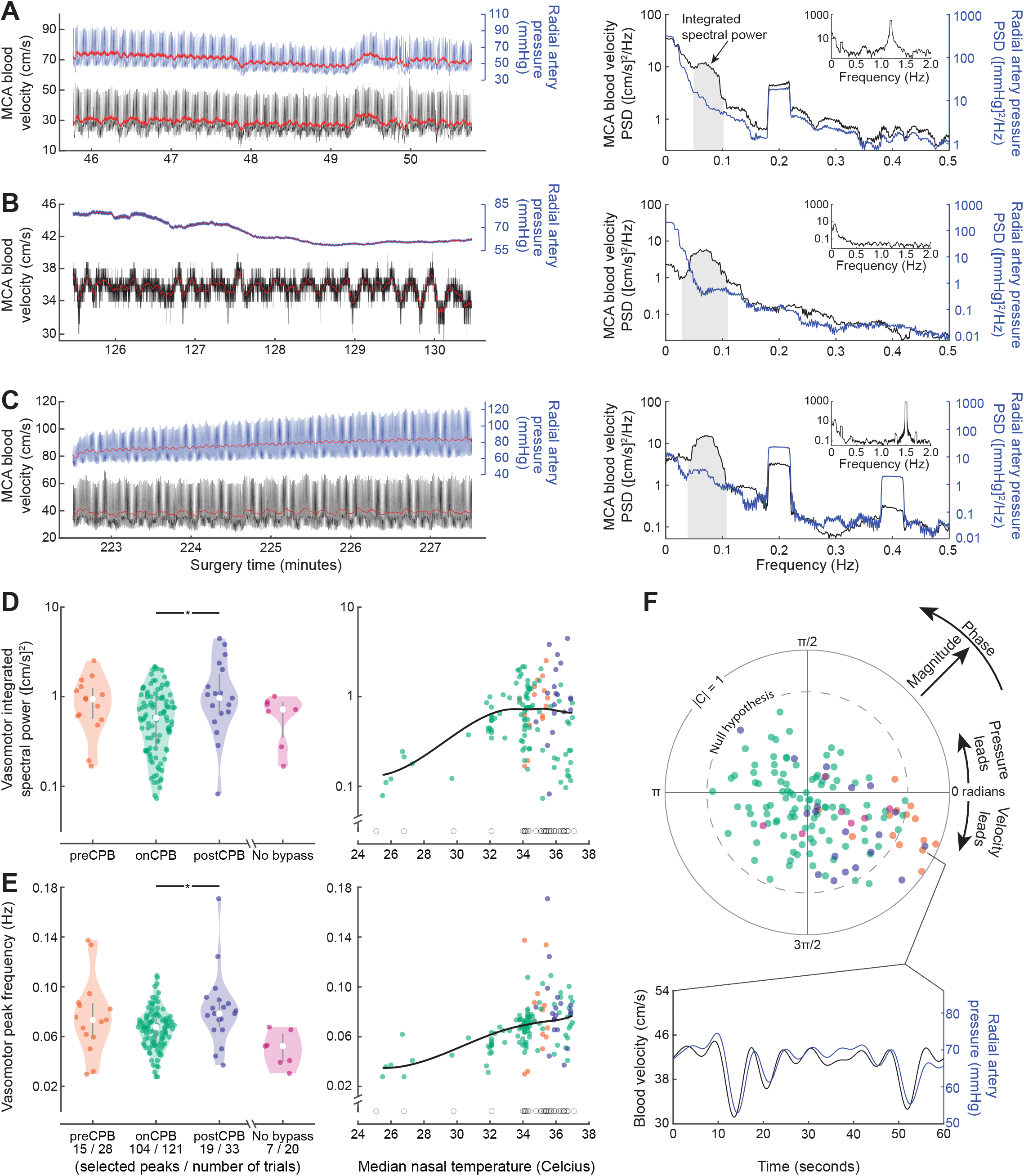
Spectral and cross-spectral analysis of MCA blood velocity and ABP signals at different stages of CPB. **A-C**, Example 5-minute recordings and PSD from a single patient pre-CPB (**A**), on-CPB (**B**), and post-CPB (**C**), with two-second moving mean traces overlaid in red. Insets: MCA blood velocity PSD, with expanded frequency range showing heart-beat rhythms. **D**, MCA blood velocity integrated spectral power, by surgical period and versus temperature. Median, 25^th^, and 75^th^percentiles are overlaid on violin plots and significant differences are indicated by * (p < 0.05). Trials with no vasomotor peak are plotted as open circles. The kernel-based 50^th^percentile estimate is overlaid in black. **E**, Vasomotor peak frequency by surgical period and versus temperature. **F**, Coherence between MCA blood velocity and concurrently measured ABP. Two trials with |C| = 0.44 and |C| = 0.14 and respective null hypothesis thresholds 0.79 and 0.76 were not plotted. Bandpass filtered signals (0.02 – 0.18 Hz) are shown for one example trial.

Low frequency oscillations were identified using a blind, manual peak-selection procedure. Plots of PSD were presented, in random order and without labels as to subject or experimental period, from a list of all pressure and velocity recordings. Further, PSD values were omitted to ensure that peak selection was blind to the identity of each spectrum. Spectra with peaks between 0.02 and 0.2 Hz and prominence above the spectral background were identified, and the frequency-endpoints of these peaks were selected. Vasomotor power was calculated by integrating the PSD between the selected frequency-endpoints and the vasomotor peak frequency was calculated as the midpoint of the frequency-endpoints. Recordings that did not have a definable vasomotor peak were excluded from statistical analyses and plotted as open circles.

### Statistics

Statistical analyses were performed in R (version 4.5.2, 2025); results with p < 0.05 were considered statistically significant unless otherwise noted. Group analyses were performed under the assumption that all recordings were independent. Shapiro-Wilk tests indicated that the on-CPB vasomotor power and post-CPB peak frequency groups deviated significantly from normality (p = 0.014 and 0.011 respectively). Kruskal-Wallis tests followed by two-sided Wilcoxon rank-sum tests with Bonferroni correction were used to identify differences between data grouped by surgical period (**Figure 2D,E**). The no bypass group was excluded from this analysis. Spearman’s rank correlation coefficients (r_s_) were calculated for both vasomotor power versus temperature and vasomotor frequency versus temperature joint distributions, after Henze-Zirkler tests^27^rejected the null hypothesis that the distributions were multivariate normal (p < 0.001 in both cases). For these datasets, 50^th^percentile estimates were found using quantile regression with second order Epanechnikov kernels and 1.25°Celsius x-bandwidths^28^. Estimates were fit with splines using MATLAB’s “spap2” function before plotting. For each trial with a peak in the PSD of the blood velocity, the magnitude and phase of the spectral coherence at the vasomotor peak frequency was displayed on a polar plot (**Figure 2F**). To determine which of these trials exhibited significant coherence, we calculated the significance level under the assumption of Gaussian statistics, i.e., 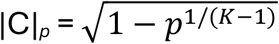 for *p* = 0.001 and *K* tapers. This assumption was validated by calculating |C| for randomized surrogate data drawn from the time series under investigation. Under the null hypothesis of zero coherence, |C| exceeds |C|_*p*_ in *p* x 100 % of trials.

## Results

In this study we used transcranial Doppler ultrasound (TCD), a non-invasive technique that measures the velocity of blood flow in the major intracranial arteries, during CPB surgery. Its sampling frequency, i.e., greater than 100 Hz, makes it suitable for assessing dynamic blood flow changes in real time^1^. TCD has been increasingly used during CPB to monitor cerebral blood flow dynamics in real time with the middle cerebral artery (MCA) as a common target, based on accessibility and clinical relevance (**insert, Figure 1A**). Arterial blood pressure (ABP) measured at the radial artery served as a control for residual Mayer wave-like blood pressure oscillations that may persist during CPB.

Twenty-two patients were approached between 2020 and 2023. Suitable TCD data was retrieved from 16 patients (11 males and 5 females) with an average age of 66 (36-85) years. Data on-CPB was retrieved from 14 patients while the other two only had traces off-CPB (**Table 1**). Simultaneous ABP and TCD recordings were collected during all surgical timepoints and were included in the analysis if the signals appeared to be stable and of good quality, following visual inspection (**Figure 1B-D**).

**Table 1:**
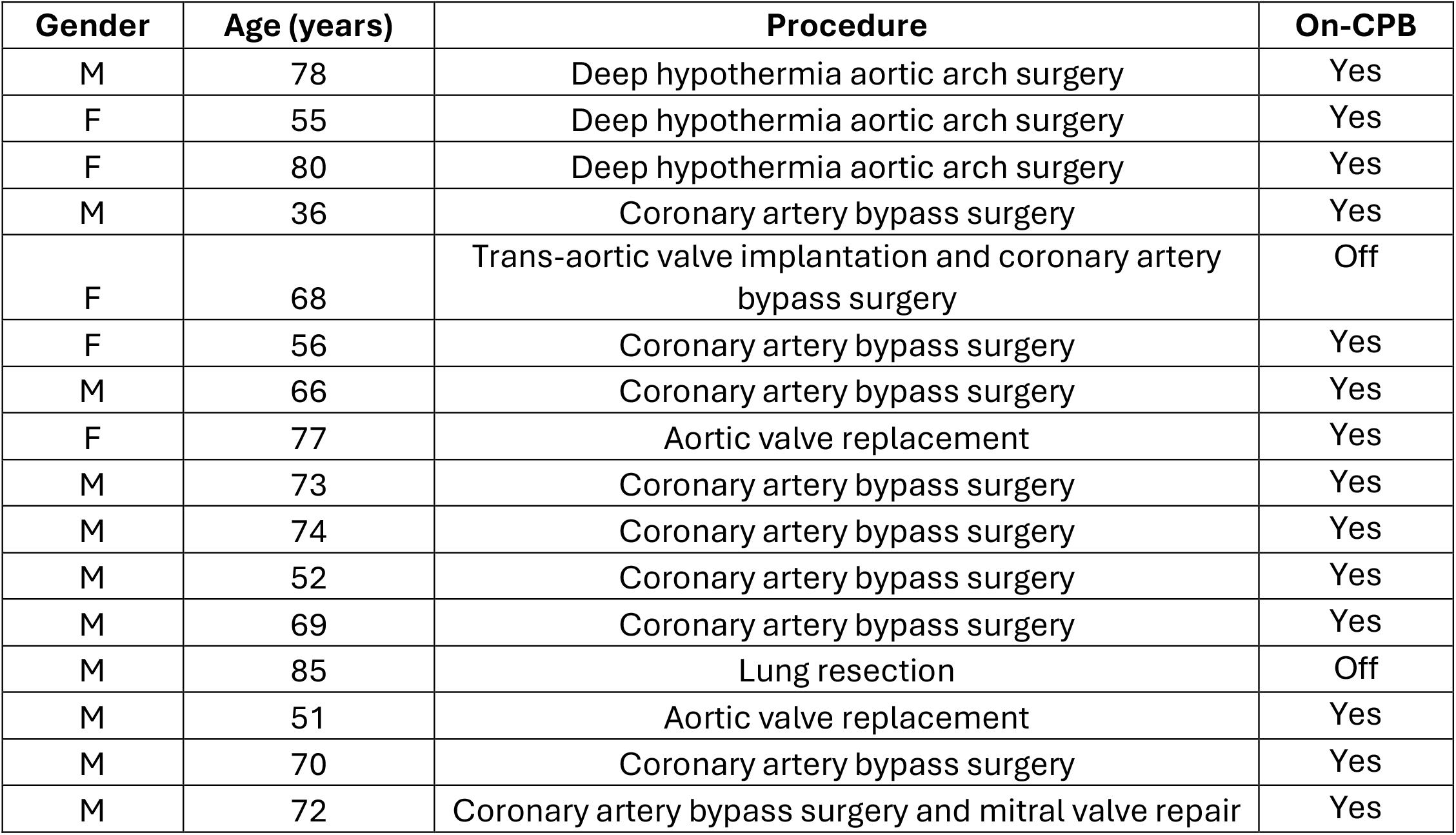
Participant demographics and procedures.

Each of the 16 subjects in the study displayed low frequency MCA blood velocity oscillations near 0.1 Hz. These oscillations were present during pre-CPB, on-CPB, and post-CPB periods and produced a characteristic peak in the blood velocity PSD (**Figure 2A-C**). Across subjects, 54 % (15 of 28) of trials exhibited a low-frequency peak during pre-CPB. During normal conditions, we expect that incoherent cardiac activity may obscure intrinsic VSMC oscillations. Indeed, when the heart was isolated from the circulation, the occurrence of these low frequency peaks in the PSD increased to 86 % (104 of 121 trials) (**Figure 2D, E**). The observed oscillations were statistically unchanged from pre-CPB to on-CPB in terms of both vasomotor power and peak frequency (**Figure 2D, E**; p = 0.19 and 0.56 respectively, two-sided Wilcoxon rank-sum tests). Vasomotor power and peak frequency differed significantly between on-CPB and post-CPB periods (p = 0.02 and 0.01 respectively, two-sided Wilcoxon rank-sum tests), although the difference in median values in each case were small (0.19 cm^2^/s^2^and 0.01 Hz respectively).

Low frequency oscillations were less common in arterial pressure recordings. The PSD for ABP were generally featureless in the ∼ 0.1 Hz range (**Figure 2A-C**), and definable peaks were present in only 23 of 202 trials across subjects and surgical periods. However, it is still possible that the observed ∼ 0.1 Hz oscillations in velocity were influenced by ABP activity in the same frequency band. For this reason, we calculated the coherence between the two signals as a measure of their mutual linear influence. We found that 92 % (134 of 145) of trials with blood velocity vasomotor peaks had insignificant magnitude coherence between blood velocity and ABP (**Figure 2F**). We conclude that the observed persistent MCA blood velocity oscillations are not dependent on oscillations in blood pressure and are caused by intrinsic vasomotion.

Vasomotor oscillations vary with temperature in onset, frequency, and amplitude^5, 29^. We found that vasomotor power varied with temperature measured at the nasopharynx, but that the two were not significantly correlated over a range of hypothermic to normal body temperatures (**Figure 2D**; Spearman’s rank correlation r_s_= 0.08, p = 0.34). Peak frequency was significantly correlated with temperature over the same range (**Figure 2E**; Spearman’s rank correlation r_s_= 0.51, p = 6×10^-10^), consistent with animal studies which show that vasomotor frequencies increase with temperature^29, 30^. Further, recent functional magnetic resonance imaging (fMRI) studies indicate that VSMC activity may be slower in humans than in rodents^31^. Our results across subjects and trials yield a peak vasomotor frequency of 0.07 ± 0.02 Hz (mean ± SD), lower than the peak frequency of 0.09 ± 0.04 Hz (mean ± SD) measured using a VSMC calcium indicator in mice^19^.

The few trials with significant coherence between blood velocity and ABP, i.e., 11 of 145, exhibited a consistent temporal relationship, where MCA blood velocity preceded ABP by 1.1 ± 0.8 s (mean ± SD, **Figure 2F**). This temporal relationship is consistent with cerebral autoregulation as characterized previously^4, 32, 33^. These trials occurred primarily during pre-CPB, i.e., 8 of 15 pre-CPB trials with vasomotor peaks had significant magnitude coherence, indicating that this autoregulatory process remained intact, at least during the initial surgery portion.

## Discussion

Oscillations near ∼ 0.1 Hz are a common feature of TCD recordings at rest^4, 34^. Interventions such as periodically inflated thigh-cuffs, lower-body negative pressure oscillations, and periodic sit-stand maneuvers have been used to enhance systemic blood pressure oscillations near 0.1 Hz and characterize dynamic cerebral autoregulation (reviewed by Claassen et al.^12^). Here we employed the opposite intervention, i.e., elimination of ∼ 0.1 Hz blood pressure oscillations that are subscribed to cardiac Mayer waves, to assess the nature of cerebral artery dynamics in the absence of systemic input. We eliminated coherent blood velocity-ABP oscillations in all but a small subset of trials (**Figure 2F**). When ABP oscillations were eliminated, velocity oscillations remained present in TCD recordings from the MCA (**Figures 1A,C and 2B**), indicating that ∼ 0.1 Hz cerebral vasomotion is not driven exclusively by Mayer waves. In total, our findings indicate that cerebral arteries undergo intrinsic vasomotion.

Limitations of this study are twofold. *First*, measurements were performed on subjects during anaesthesia and in some cases during periods of hypothermia. While Mayer waves have been classified during various non-surgical conditions^3^, further work is needed to delineate the intrinsic contribution of vasomotion to blood flow dynamics during awake conditions. To this end, recent work identified significant correlation between ∼ 0.1 Hz oscillations in peripheral organs, including the heart, and a spatially uniform component of the blood oxygen level-dependent (BOLD) signal^13, 35^. Our data does not dispute this claim, which is based on a correlative connection. Brain vasomotor oscillations can be independent of systemic oscillations in the body, yet the two can phase-lock if there is a sufficiently strong interaction^36^. *Second*, the precise vascular origin of ABP-independent MCA blood velocity oscillations in our study is unknown. The MCA diameter is generally accepted to be constant within normal blood gas and blood pressure ranges, supporting our assumption that TCD oscillations reflect pial and parenchymal vasomotion^37^. In rodents, arteriole vasomotion is synchronized across large areas of the brain^19^; the consequent resistance modulation would presumably cause changes in MCA flow velocity. However, the degree of vasomotion synchronization in humans is unknown. The opposing viewpoint, that TCD oscillations are the consequence of local MCA vasomotion, cannot be ruled out^38^.

Our study demonstrates vasomotion in humans when blood pressure is tightly controlled. The confounding impact of slow blood pressure oscillations - Mayer waves - on blood flow was thus eliminated. This provides unprecedented confirmation for the presence of intrinsic, rhythmic cerebrovascular activity, which is likely to contribute to perivascular clearance of amyloid and other deleterious peptides^14, 15^. This setup can be a potential platform for further observations and subsequent development of a non-invasive TCD based imaging tool that could lead to early detection of failure of perivascular clearance before the onset of dementia.

## Acknowledgements

We thank Patrick Drew, Jonathan Polimeni and Andy Shih for discussions. Funded by University of Southampton Hospital Trust, the Leducq Foundation and the Leducq Foundation for Cardiovascular Research (23CVD03), the NIH BRAIN Initiative grant U19 NS123717 and NINDS grant R35 NS097265 to D.K. The funders had no role in study design, data collection and analysis, decision to publish or preparation of the manuscript.

## Author contributions

AA and ROC conceived the study and obtained Integrated Research Application System approval for the use of human participants; AA performed surgery; AAB acquired physiological data; JD, DK and DMS analyzed the data; AA, JD, DK and ROC wrote the manuscript; and DK and ROC obtained funding.

## Statements and Declarations

### Ethical considerations

This observational study received ethical approval from the Integrated Research Application System (approval #264726) on 04/01/2019. All patient information was de-identified and patient data will not be shared with third parties. The protocol has been agreed and accepted and the Chief Investigator agreed to conduct the study in compliance with the approved protocol and will adhere to the principles outlined in the Declaration of Helsinki.

### Consent to participate

Written informed consent for inclusion in this research was obtained from the patients prior to surgery.

### Data and code availability

Raw data in NWB format will be deposited on DANDI upon acceptance. Code used for analysis is publicly available at https://github.com/JDuckworth1/HumanCBF.

## References

1. Willie CK, Tzeng Y, Fisher JA, et al. Integrative regulation of human brain blood flow. Journal of Physiology 2014; 592: 841–859.

2. Hamner JW, Cohen MA, Seiji M, et al. Spectral indices of human cerebral blood flow control: responses to augmented blood pressure oscillations. Journal of Physiology 2004; 559: 965–973.

3. Julien C. The enigma of Mayer waves: facts and models. Cardiovascular Research 2006; 70: 12–21.

4. Diehl RR, Linden D, Lücke D, et al. Spontaneous blood pressure oscillations and cerebral autoregulation. Clinical Autonomic Research 1998; 8: 7–12.

5. Intaglietta M. Vasomotion and flowmotion: Physiological mechanisms and clinical evidence. Reviews in Vascular Medicine 1990; 1: 101–112.

6. Aalkjær C, Boedtkjer D and Matchkov V. Vasomotion – what is currently thought? Acta Physiologiae 2011; 202: 253–269.

7. Rayshubskiy A, Wojtasiewicz TJ, Mikell CB, et al. Direct, intraoperative observation of∼ 0.1 Hz hemodynamic oscillations in awake human cortex: Implications for fMRI. NeuroImage 2013; 87: 323–331.

8. Lecrux C and Hamel E. Neuronal networks and mediators of cortical neurovascular coupling responses in normal and altered brain states. Philosophical Transactions of the Royal Society of London B 2016; 371: e20150350.

9. Iadecola C. The neurovascular unit coming of age: a journey through neurovascular coupling in health and disease. Neuron 2017; 96: 15–42.

10. Mateo C, Knutsen PM, Tsai PS, et al. Entrainment of arteriole vasomotor fluctuations by neural activity is a basis of blood oxygen level dependent “resting state” connectivity. Neuron 2017; 96: 936–948.

11. Claassen JAHR, Levine BD and Zhang R. Dynamic cerebral autoregulation during repeated squat-stand maneuvers. Journal of Applied physiology 2009; 106: 153–160.

12. Claassen J, Thijssen DHJ, Panerai RB, et al. Regulation of cerebral blood flow in humans: physiology and clinical implications of autoregulation. Physiological Reviews 2021; 101: 1487–1559.

13. Bolt T, Wang S, Nomi JS, et al. Autonomic physiological coupling of the global fMRI signal. Nature Neuroscience 2024; 28: 1327–1335.

14. Aldea R, Weller RO, Wilcock DM, et al. Cerebrovascular smooth muscle cells as the drivers of intramural periarterial drainage of the brain. Frontiers in Aging Neuroscience 2019; 11: e1.

15. van Veluw SJ, Hou SS, Calvo-Rodriguez M, et al. Vasomotion as a driving force for paravascular clearance in the awake mouse brain. Neuron 2020; 105: 549–561.

16. Hladky SB and Barrand MA. The glymphatic hypothesis: The theory and the evidence. Fluids and Barriers of the CNS 2022; 19: e282.

17. Gan Y, Holstein-Rønsbo S, Nedergaard M, et al. Perivascular pumping of cerebrospinal fluid in the brain with a valve mechanism. Journal of the Royal Society Interface 2023; 20: e20230288.

18. Munting LP, Bonnar O, Kozberg MG, et al. Spontaneous vasomotion propagates along pial arterioles in the awake mouse brain like stimulus-evoked vascular reactivity. Journal of Cerebral Blood Flow & Metabolism 2023; 43: 1752–1763.

19. Broggini T, Duckworth J, Ji X, et al. Long-wavelength traveling waves of vasomotion modulate the perfusion of cortex. Neuron 2024; 112: 2349–2367.

20. Kedarasetti RT, Drew PJ and Costanzo F. Arterial vasodilation drives convective fluid flow in the brain: a poroelastic model. Fluids and Barriers of the CNS 2022; 19: e34.

21. Nedergaard M and Goldman SA. Science. Glymphatic failure as a final common pathway to dementia 2020; 370: 50–56.

22. Bumstead JR, Bauer AQ, Wright PW, et al. Cerebral functional connectivity and Mayer waves in mice: Phenomena and separability. Journal of Cerebral Blood Flow and Metabolism 2016; 37: 471–484.

23. Podgoreanu MV, Stout RG, El-Moalem HE, et al. Synchronous rhythmical vasomotion in the human cutaneous microvasculature during nonpulsatile cardiopulmonary bypass. Anesthesiology 2002; 97: 1110–1117.

24. Boettcher W, Merkle F and Heinz-Hermann W. History of extracorporeal circulation: the conceptional and developmental period. Journal of Extra Corporeal Technology 2003; 35: 172–183.

25. Swain JA, McDonald TJJ, Griffith PK, et al. Low-flow hypothermic cardiopulmonary bypass protects the brain. Journal of Thoracic and Cardiovascular Surgery 1991; 102: 76–84.

26. Percival DB and Walden AT. Spectral Analysis for Physical Applications: Multitaper and Conventional Univariate Techniques. Cambridge: Cambridge University Press, 1993.

27. Korkmaz S, Goksuluk D and Zararsiz G. MVN: An R package for assessing multivariate normality. The R Journal 2014; 6: 151–162.

28. Hayfield T and Racine JS. Nonparametric econometrics: the np package. Journal of Statistical Software 2008; 27.

29. Setchell BP, Bergh A, Widmark A, et al. Effect of testicular temperature on vasomotion and blood flow. International Journal of Andrology 1995; 18: 120–126.

30. Bouskela E. Vasomotion frequency and amplitude related to intraluminal pressure and temperature in the wing of the intact, unanesthetized bat. Microvascular Research 1989; 37: 339–351.

31. Lambers H, Segeroth M, Albers F, et al. NeuroImage. A cortical rat hemodynamic response function for improved detection of BOLD activation under common experimental conditions 2020; 208: e116446.

32. Diehl RR, Linden D, Lücke D, et al. Phase relationship between cerebral blood flow velocity and blood pressure: a clinical test of autoregulation. Stroke 1995; 26: 1801–1804.

33. Hughson RL, Edwards MR, O’Leary DD, et al. Critical analysis of cerebrovascular autoregulation during repeated head-up tilt. Stroke 2001; 32: 403–2408.

34. Diehl RR, Diehl B, Sitzer M, et al. Spontaneous oscillations in cerebral blood flow velocity in normal humans and in patients with carotid artery disease. Neuroscience Letters 1991; 127: 5–8.

35. Özbay PS, Chang C, Picchioni D, et al. Sympathetic activity contributes to the fMRI signal. Communications Biology 2019; 2: e421.

36. Kuramoto Y. Chemical Oscillations, Waves and Turbulence. New York: Springer Verlag, 1984.

37. Aaslid R, Lindegaard KF, Sorteberg W, et al. Cerebral autoregulation dynamics in humans. Stroke 1989; 20: 45–52.

38. Hoiland RL and Ainslie PN. CrossTalk proposal: The middle cerebral artery diameter does change during alterations in arterial blood gases and blood pressure. Journal of Physiology 2016; 594: 4073–4075.

